# Shifts in outcrossing rates and changes to floral traits are associated with the evolution of herbicide resistance in the common morning glory

**DOI:** 10.1101/061689

**Authors:** Adam Kuester, Eva Fall, Shu-Mei Chang, Regina S. Baucom

## Abstract

Human-mediated selection can strongly influence the evolutionary response of natural organisms within ecological timescales. But what traits allow for, or even facilitate, adaptation to the strong selection humans impose on natural systems? Using a combination of lab and greenhouse studies of 32 natural populations of the common agricultural weed, *Ipomoea purpurea*, we show that herbicide resistant populations self-fertilize more than susceptible populations. We likewise show that anther-stigma distance, a floral trait associated with self-fertilization in this species, exhibits a non-linear relationship with resistance such that the most and least resistant populations exhibit lower anther-stigma separation compared to populations with moderate levels of resistance. Overall, our results extend the general finding that plant mating can be impacted by human-mediated agents of selection to that of the extreme selection of the agricultural system. This work highlights the influence of human-mediated selection on rapid responses of natural populations that can lead to unexpected long-term evolutionary consequences.

**Statement of authorship:** AK collected seeds, performed experiments, analyzed data and wrote the paper; EF collected data; SMC collected data and contributed to the manuscript; RSB designed the study, performed the analyses, and wrote the paper. All authors discussed the results and commented on the manuscript.

**Data accessibility:** Primary data used in these analyses will be made available in the public github repository https://github.com/rsbaucom/MatingSystem2015, which can be anonymously accessed. Upon acceptance, data will be made available through the Dryad public repository.

## Introduction

Pesticides are used world-wide to protect agricultural crops from the damaging effects of insects, fungi and weeds (Enserink *et al.* 2013) and are considered vital for maintaining the world’s food supply (Lamberth *et al.* 2013). Recently we have begun to recognize that their use can negatively impact the reproduction and mating patterns of natural organisms such as bees (Williams *et al.* 2015), amphibians (Rohr & McCoy 2010), and plants (Pline *et al.* 2003; Thomas *et al.* 2004; Baucom *et al.* 2008; Londo *et al.* 2014) and can have long term evolutionary consequences on these non-target species. In the US, 40% of the pesticides applied across the 400 million acres of cropland are herbicides (*US EPA* 2011), which impose extreme selection on naturally occurring agricultural weeds (Jasieniuk *et al.* 1996; Vigueira *et al.* 2013). Strikingly, while herbicide resistance has evolved in over 200 plant species worldwide (Heap 2014), the impact on correlated, non-target traits has been largely unexplored.

The plant mating system, or the relative rate of outcrossing versus selfing, is a labile trait (Barrett 1998; Karron *et al.* 2012) that is influenced by human impacts on natural ecosystems (Eckert *et al.* 2010). Habitat fragmentation by deforestation, for example, reduces the outcrossing rate of many forest tree species (Aguilar *et al.* 2006; Eckert *et al.* 2010). Further, plants tolerant to heavy-metal contaminated soils exhibit higher rates of autonomous self-pollination than non-adapted biotypes (Antonovics *et al.* 1971), strongly suggesting that the mating system is either influenced by or concomitantly evolves in response to heavy metal exposure (Antonovics 1968; Antonovics *et al.* 1971). We predict that the mating system of an agricultural weed should co-vary with the level of herbicide resistance in nature, for two main reasons. First, the reproductive assurance hypothesis (Baker 1955, 1974; Goodwillie *et al.* 2005; Pannell *et al.* 2015) would predict resistant individuals, in a mate-limited population following herbicide application, are more likely to produce progeny if they are also highly self-pollinating rather than outcrossing. Second, resistant types that self-pollinate would effectively reduce the influx of non-adapted, susceptible alleles – otherwise known as the ‘prevention of gene flow’ hypothesis (Antonovics 1968). Both hypotheses predict that herbicide resistant individuals should self to a higher degree than non-resistant individuals. Interestingly, while many herbicide resistant weed species are reported to be predominantly selfing (Jasieniuk *et al.* 1996), there are no investigations, to our knowledge, that examine the potential for co-variation between mating system and herbicide resistance in nature, a finding that would indicate the mating system may co-evolve when populations respond to the strong selection imparted by herbicides.

*Ipomoea purpurea,* an annual weed of agricultural fields and disturbed sites in the southeastern and Midwest US, is a model for examining persistence in stressful and competitive environments (Baucom *et al.* 2011; Chaney & Baucom 2014). As such the species is a particularly relevant candidate for studying how mating systems may evolve under regimes of human-mediated selection. Populations of this agricultural weed have been exposed consistently to the application of glyphosate, the active ingredient in the herbicide RoundUp, since the late 1990’s given the widespread adoption of RoundUp Ready crops in the US (NASS 2015). Populations vary for the level of resistance to glyphosate across its North American range; while some populations of *I. purpurea* exhibit 100% survival following application of the field-dose of the herbicide, other populations exhibit high susceptibility (Fig. 1a) (Kuester *et al.* 2015a). In addition, individuals of this mixed-mating, hermaphroditic species (Ennos 1981; Brown & Clegg 1984) with a smaller anther-stigma distance (ASD, the distance between the tallest anther and the stigma; Fig. 1b) self-pollinate more often than individuals with a larger ASD (Ennos 1981; Chang & Rausher 1999). Because both anther-stigma distance and glyphosate resistance in this species are heritable and respond to selection (Ennos 1981; Chang & Rausher 1999; Debban *et al.* 2015) this common agricultural weed provides a unique opportunity to examine the potential that mating systems and associated reproductive traits evolve in response to extreme regimes of selection imposed by herbicide.

**Figure 1.**
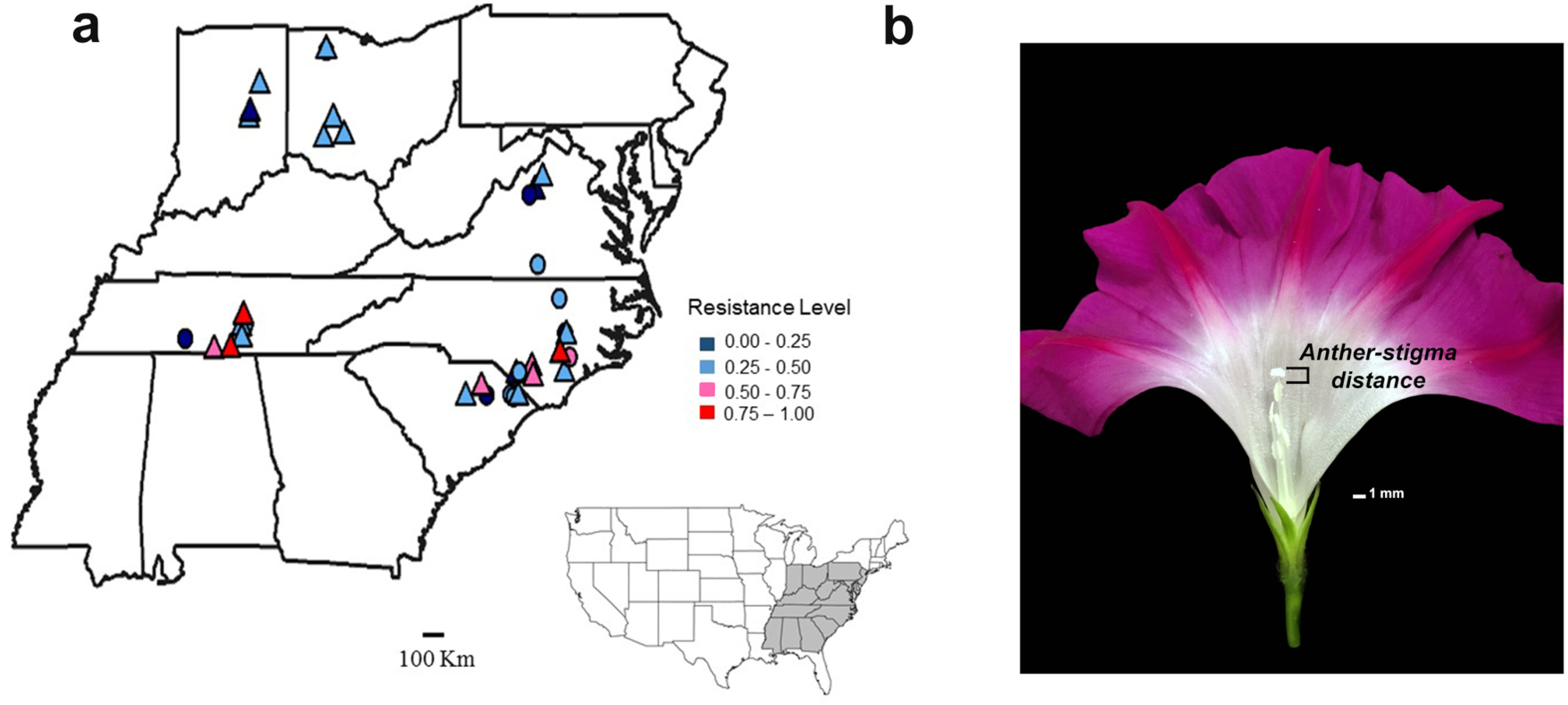
Map of populations sampled within the USA, **a**, and image of anther-stigma distance in *I. purpurea,* **b**. Floral morphology was measured for all populations (N = 32) whereas the mating system was estimated for populations indicated by triangles (N = 24). The color indicates the resistance level for each population based on proportion survival following application of 1.7 kg ai/ha of herbicide, which is slightly higher than the recommended field dose of herbicide (from Kuester *et al.* 2015a). Sites were sampled at least 5 km apart.

Here, we determine if the mating system of *Ipomoea purpurea* co-varies with herbicide resistance, and if reproductive traits associated with self-fertilization are likewise influenced by resistance status. We previously estimated the percent survival of populations following herbicide application using a replicated dose-response greenhouse experiment with individuals sampled as seed from the field in 2012. We used these population-level estimates of survival as each population’s resistance level (Fig. 1a) (Kuester *et al.* 2015a), and determined if molecular-marker based estimates of the mating system and measurements of floral morphology co-varied with the level of herbicide resistance in natural populations. Further, our experimental populations were sampled twice, once in 2003 and again in 2012 from the same location (Fig. 1a), allowing us to examine the hypothesis that floral traits have changed in these populations over time. We predict that populations with a high level of glyphosate resistance should exhibit evidence of reduced outcrossing and reductions in the anther-stigma distance – either of which would indicate that plant reproductive traits exhibit correlated evolution in response to strong selection from herbicide application.

## Materials and Methods

### Mating system estimates

We performed a progeny array analysis to estimate the mating system of 24 populations located in the southeastern and Midwestern US (indicated by triangles in Fig. 1a). These populations are part of a previous study in which we screened for glyphosate resistance, defined as the proportion each population that survived the application of 1.7 kg a.i./ha RoundUp (Kuester *et al.* 2015a), a rate that is slightly higher than the current recommended field dose. One seed randomly selected from an average of 11 fruits per maternal line from an average of 19 maternal lines per population were used to estimate mating system parameters (see Table S1 in Supporting Information for numbers of maternal lines and progeny per population). Maternal plants were sampled at least two meters apart within each agricultural field. DNA was extracted from seedling cotyledon tissues using a CTAB method (T Culley, pers. comm.). In total, 4798 progeny were genotyped with fifteen previously described microsatellite loci for maternal-line estimates of the mating system (see Kuester *et al.* 2015a) for specific details of the PCR conditions). All sampled genotypes were analyzed using Applied Biosystems PeakScanner 1.0 analytical software (Carlsbad, CA) with a PP (Primer Peaks adjustment) sizing default, and scoring was double-checked manually for errors in a random sub-sample of 200 individuals. We examined our ability to assign parentage using Cervus (Kalinowski *et al.* 2007) and determined population-level estimates of genetic diversity and inbreeding using GenalEx (Peakall & Smouse 2012). We estimated mating system parameters using BORICE (Koelling *et al.* 2012), which is a Bayesian method to estimate the family-level outcrossing rate (t) and maternal line inbreeding coefficients (F) (Koelling *et al.* 2012). BORICE is reported to perform well when either family sizes are small or maternal genotypes are unavailable (Koelling *et al.* 2012). We used the following default parameters when estimating mating system parameters: 1 million iterations and 99,999 burn-in steps, an outcrossing rate tuning parameter of 0.05, an allele frequency tuning parameter of 0.1, and an initial population outcrossing rate of 0.5. We examined the possibility that null alleles influenced our mating system estimates by re-running all analyses after excluding 4 loci that potentially exhibited null alleles (i.e., loci with ~25% null alleles: IP18, IP1, IP26 and IP42) as indicated by MicroChecker (Van Oosterhout et al. 2004). We found no evidence that null alleles impacted estimates across populations (correlation between outcrossing rates for all loci and 4 loci removed: r = 0.94, P < 0.001) and thus report results using all 15 loci.

### Floral phenotypes

We performed a replicated greenhouse experiment to determine if floral morphology varied according to resistance level and to determine if floral traits differed between two sampling years (2003 and 2012). Seeds were sampled from maternal plants from 32 randomly chosen populations located in the Southeast and Midwest US in the fall of 2012 (all populations in Fig. 1a); fifteen of these populations had been previously sampled in 2003 (see Table S2). We planted seeds from between 1-29 maternal lines (average=12.74, median=13; see Table S2 for number of individuals) from each population and each sampling year in 4-inch pots in a completely randomized design at the Matthaei Botanical Gardens at the University of Michigan (Ann Arbor, MI). To increase overall sample size, a second replicate experiment was started two weeks later in the greenhouses for a total of 640 experimental plants. Once plants began flowering, we measured the height of the pistil (cm) and the tallest stamen (cm) to estimate anther-stigma distance (ASD) of an average of 5.5 flowers per plant across 17 sampling dates. An average of 2 flowers were measured from each plant each sampling date; measurements taken on multiple flowers per plant per sampling date were averaged prior to analysis. Over the course of the experiment, we measured 3569 flowers from 622 experimental plants. Of the overall 32 populations sampled in 2012 for floral morphology estimates, 23 were likewise represented in the mating system analyses, presented above.

### Statistical analyses

To determine if mating system parameters (the outcrossing rate (t), maternal inbreeding coefficient (F)) of *I. purpurea* co-varied with resistance, we performed linear regressions using the lm function in R v 3.1.1 (R Core Team 2013) in which each population’s mating system estimate was used as the dependent variable with the level of resistance (proportion survival at 1.7 kg a.i./ha) as an independent variable. We included population latitude in preliminary models as an independent variable since previous work (Kuester *et al.* 2015a) indicated a weak trend between resistance and latitude; however, this effect was removed from final models as it was never significant nor did it influence the relationship between mating system and resistance. Preliminary analyses indicated the presence of a nonlinear relationship between the outcrossing rate and resistance, and as such we included a quadratic term in a separate regression for both mating system parameters. Further, because a plot of the outcrossing rate and the quadratic resistance term exhibited a non-linear but not completely convex relationship, we performed a piece-wise regression to examine the potential for two different linear relationships in the data. To do so, we used the segmented package of R (Muggeo 2008) with an initial approximated breakpoint (psi) of 0.40. Each mating system parameter was examined for normality by performing Shapiro-Wilk tests. Neither showed evidence of non-normality and therefore were not transformed prior to analysis.

We examined whether floral morphology differed according to the level of resistance first using populations sampled in 2012 (N = 32). To do so we performed a multivariate ANOVA with sine-transformed values of each floral phenotype (anther-stigma distance, length of the tallest stamen, and pistil height) as dependent variables in the model and the experimental replicate, resistance level of the population, and latitude of the population as fixed, independent variables. Prior to analysis, we removed the influence of sampling date (N=17) by performing a MANOVA with transformed variables and retained the residuals for testing our main effects of interest (experimental replicate, resistance level, latitude). We elected to do so because the influence of sampling date on anther-stigma distance was not one of our primary questions and we noted a highly significant influence of this effect on floral morphology. Preliminary analyses indicated a significant influence of population latitude on floral morphology and thus we elected to include this effect in the MANOVA. Further, because a scatterplot of the relationship between anther-stigma distance and resistance indicated the presence of a non-linear relationship, we included a quadratic term (resistance level^2^) in this and downstream analyses.

We next ran separate univariate analyses to determine if ASD, height of the tallest stamen and pistil height varied according to resistance level and if they have changed over sampling years using the lm function of R v 3.1.1 (R Core Team 2013). For each model, the sine-transformed floral trait of interest (ASD, height of tallest stamen, pistil height) was the dependent variable with the experimental replicate, resistance level, resistance level^2^, and population latitude as independent variables in the regression models. We again performed analyses using residuals after removing the effect of sampling date. Similar to the mating system analysis, if the non-linear resistance term proved significant, we used the segmented package to perform a piece-wise regression and examine the potential for two different linear relationships as well as the break point between slopes using an initial breakpoint of 0.40 (Muggeo 2008). To determine if floral morphology had changed between sampling years and/or if floral morphology differed between years differently across resistance levels, we performed the same analyses as above using only populations that were sampled in both 2003 and 2012 (N = 15), including a year term, a year by resistance level term, and a year by resistance level^2^ term in each analysis. In these analyses, we used resistance levels from each population each year of sampling (as reported in Kuester *et al.* 2015b).

## Results

### Mating system

The average combined exclusion probabilities across all populations was high, at greater than 99%, indicating that the fifteen microsatellite loci successfully assigned parentage (see Table S1). Thus, the power is sufficient for estimating outcrossing rates among populations. Values of the outcrossing rate varied substantially across populations (range: 0.27-0.8), with an average value (±SE) for the species of 0.50 (±0.03) (Table S3).

We uncovered a strong and striking negative linear relationship between the average family-level outcrossing rate of each population (Table S3) and the level of glyphosate resistance (β = -0.30 ±0.09; Fig. 2a); individuals from highly resistant populations self-pollinate more than individuals from less resistant populations (F = 10.70, *P* = 0.004; Table 1). We also uncovered a significant negative quadratic relationship between resistance and the outcrossing rate, suggesting that outcrossing first increases at low levels of resistance and then declines as resistance increases (F = 5.08, *P* = 0.035; Table 1). Piece-wise regression analysis indicated, however, that the slope between the outcrossing rate and resistance was positive but not significantly different from zero at low levels of resistance (β = 0.42 ±0.36, 95% CI: -0.34, 1.17) whereas the slope following the estimated break point (0.42 ±0.07) was negative and significantly different from zero (β = -0.57 ±0.14, 95% CI: -0.28, -0.86). In line with our finding of lower outcrossing in highly resistant populations, we found maternal inbreeding coefficients to increase linearly as the level of resistance increases (Fig. 2b, F = 6.05, *P* = 0.02; Table 1). We found no evidence of a nonlinear relationship between the maternal inbreeding coefficients and the level of resistance. Together, these results demonstrate that outcrossing rates were lower and maternal inbreeding coefficients higher in glyphosate resistant compared to susceptible populations.

**Table 1.** Results of separate linear and quadratic regressions testing the influence of resistance on the outcrossing rate (t) and the maternal inbreeding coefficient (F). Coefficients from the quadratic regressions included the linear term whereas coefficients from the linear regressions were determined without the quadratic term. Effects that are significant (P < 0.05) are bolded.

|  | Outcrossing rate (t) |  |  |  | Maternal inbreeding coefficient (F) |  |  |  |
| --- | --- | --- | --- | --- | --- | --- | --- | --- |
| Effect | Coefficient (SE) | Df | F | P | Coefficient (SE) | Df | F | P |
| Resistance Level | -0.30 (0.09) | 1, 22 | 10.694 | **0.004** | 0.14 (0.06) | 1, 22 | 6.051 | **0.022** |
| Resistance Level<sup>2</sup> | -0.77 (0.35) | 1, 21 | 5.084 | **0.035** | 0.13 (0.22) | 1, 21 | 0.335 | 0.569 |

**Figure 2.**
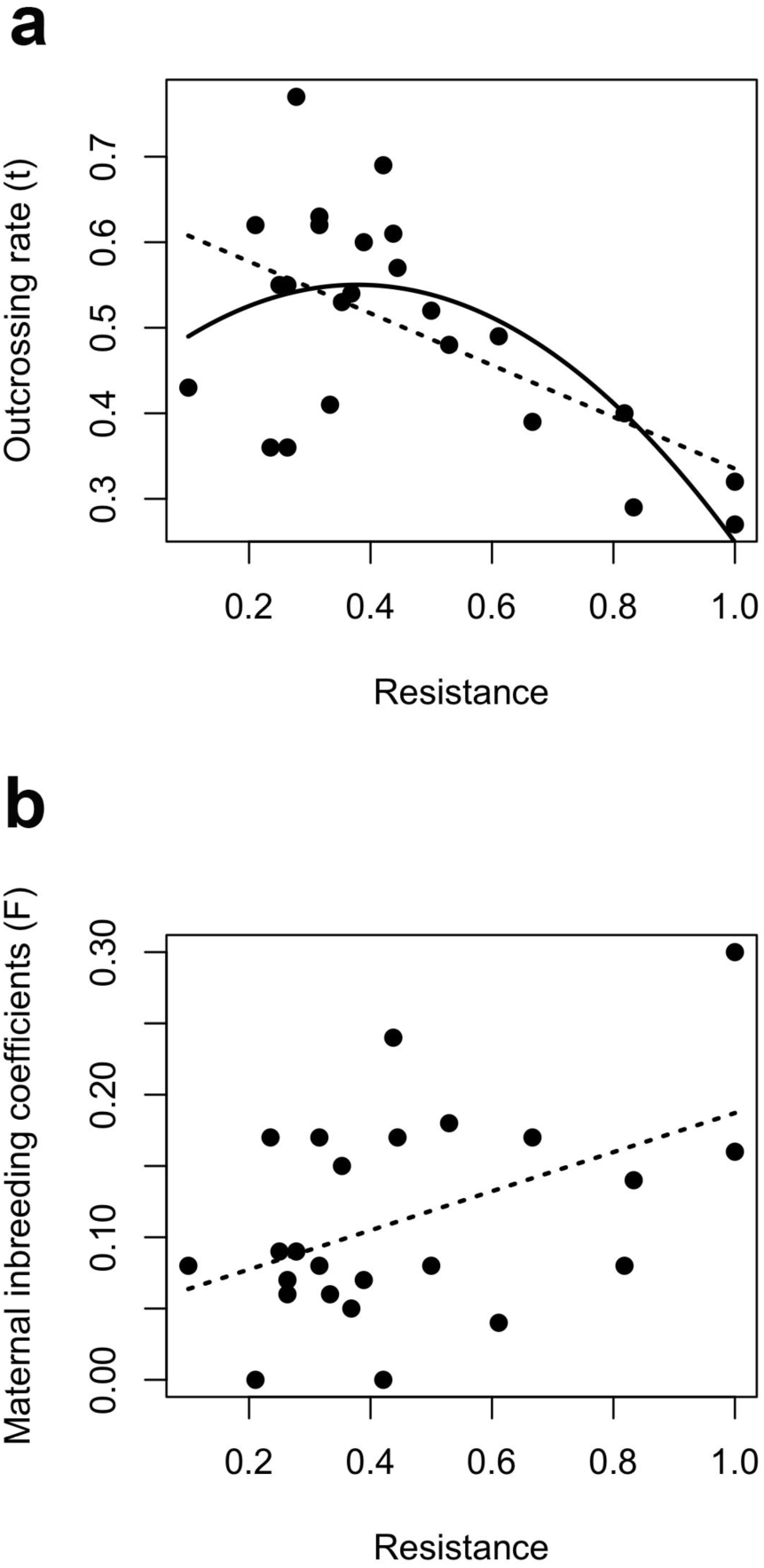
The relationship between mating system parameters and the proportion survival of each population following application of 1.7 kg ai/ha of herbicide in *I. purpurea*. **a**, outcrossing rate (t), **b**, mean inbreeding coefficient of maternal individuals (F). Significance is indicated by a regression line. A significant negative linear (dashed line) and quadratic (solid line) relationship was detected for the outcrossing rate whereas a significant positive linear relationship was uncovered for the maternal inbreeding coefficients (Table 1).

### Floral morphology

A multivariate analysis of variance indicated that floral morphology related to selfing rates (anther-stigma distance (ASD), height of the pistil and the tallest stamen) was significantly influenced by both the non-linear resistance term and the latitude of the population (resistance level^2^, Approx. F = 6.66, *P* < 0.001; latitude, Approx. F = 2.58, *P* = 0.05; Table S4). In separate univariate ANOVAs, we found a negative quadratic relationship between ASD and resistance (resistance level^2^: F = 5.70, *P* = 0.02, Table 2, Fig. 3a) and a trend for a negative quadratic relationship between resistance and pistil height (resistance level^2^: F = 2.96, *P* = 0.09, Table 2, Fig. 3c) but no quadratic relationship between resistance and stamen height (Fig. 3b). No linear relationships between resistance and the three floral traits were uncovered; however, a piece-wise regression analysis of ASD indicated a positive slope between ASD and resistance at low levels of resistance (β = 0.72 ±0.32, 95% CI: 0.06, 1.37) and a negative slope (β = -0.16 ±0.07, 95% CI: -0.01, -0.31) after the estimated breakpoint (0.31±0.05).

**Table 2.**
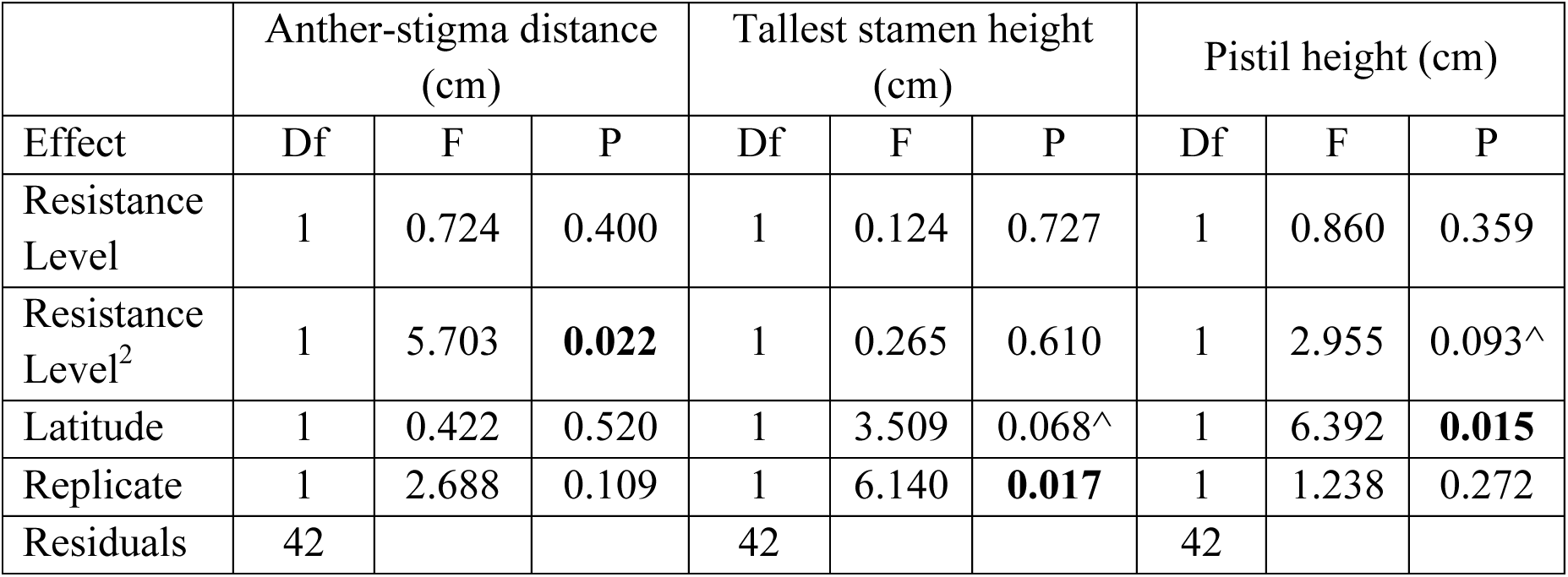
Results of separate ANOVAs testing the influence of population resistance level, resistance level^2^, population latitude and experimental replicate on anther-stigma distance (cm), pistil height (cm) and height of tallest stamen (cm). Significant effects (P < 0.05) are bolded whereas an ‘^’ indicates a trend for significance.

|  | Anther-stigma distance (cm) |  |  | Tallest stamen height (cm) |  |  | Pistil height (cm) |  |  |
| --- | --- | --- | --- | --- | --- | --- | --- | --- | --- |
| Effect | Df | F | P | Df | F | P | Df | F | P |
| Resistance Level | 1 | 0.724 | 0.400 | 1 | 0.124 | 0.727 | 1 | 0.860 | 0.359 |
| Resistance Level<sup>2</sup> | 1 | 5.703 | **0.022** | 1 | 0.265 | 0.610 | 1 | 2.955 | 0.093^ |
| Latitude | 1 | 0.422 | 0.520 | 1 | 3.509 | 0.068^ | 1 | 6.392 | **0.015** |
| Replicate | 1 | 2.688 | 0.109 | 1 | 6.140 | **0.017** | 1 | 1.238 | 0.272 |
| Residuals | 42 |  |  | 42 |  |  | 42 |  |  |

**Figure 3.**
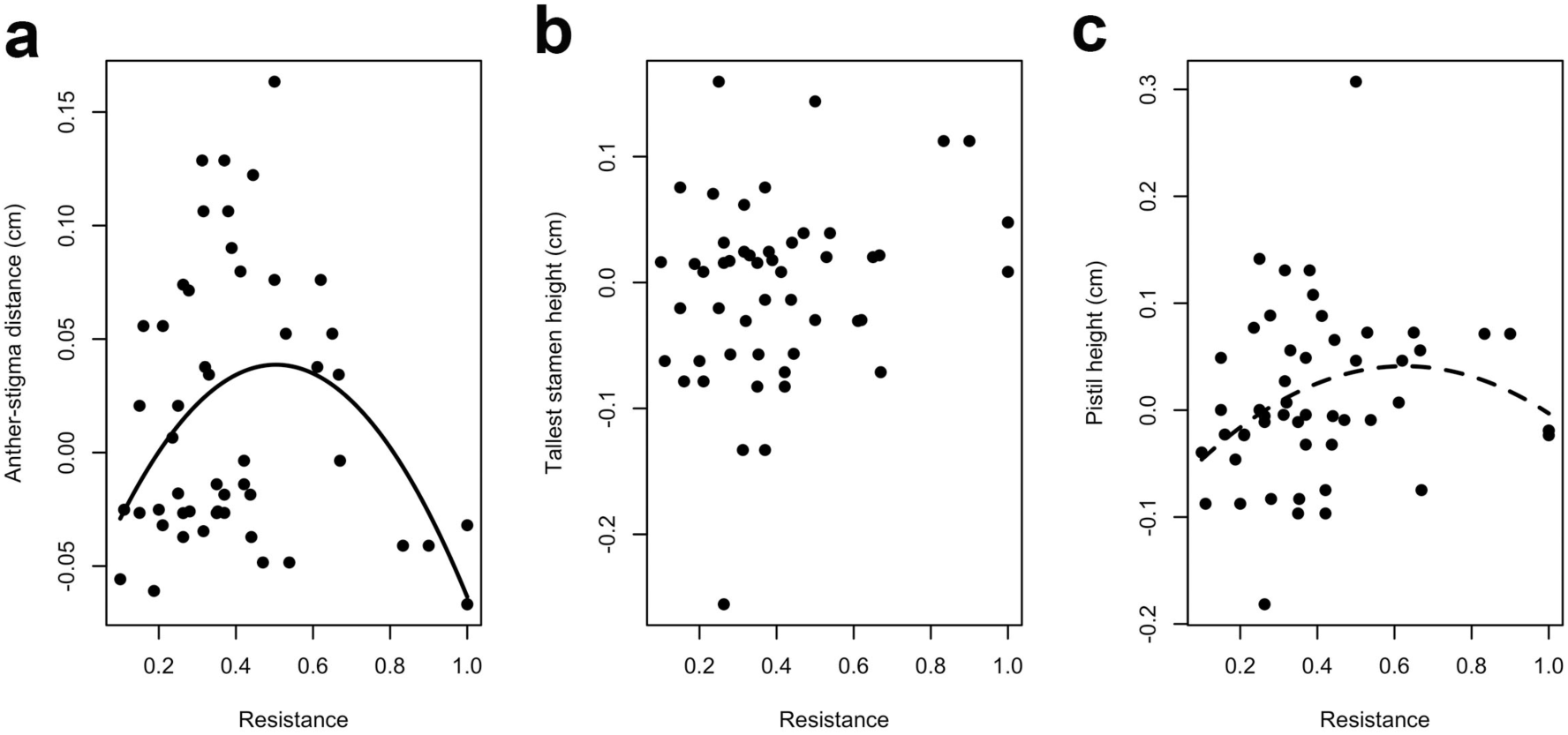
Anther-stigma distance varies non-linearly with the level of resistance. Thirty-two populations sampled as seed in 2012 were used in this analysis to determine if floral traits co-varied with variation in the level of glyphosate resistance. Shown are the residuals, averaged by population, after removing variation due to sampling date for **a**, anther-stigma distance (cm), **b**, height of the tallest stamen (cm), and **c**, pistil height (cm) according to resistance level at 1.7 kg ai/ha. A significant relationship between the trait and resistance is indicated by a solid line (P < 0.05) whereas a dashed line indicates a trend for significance P < 0.10 (see Table 2).

There was no evidence that ASD either increased or decreased between collection years across the subset of populations sampled in both 2003 and 2012 (Year effect in ANOVA: F_1,47_ = 0.464, P = 0.50). Further, although we again detected a significant quadratic relationship between ASD and resistance using this subset of populations (resistance level^2^: F_1,47_ = 5.82, P = 0.02, Fig. 4), we uncovered no evidence that this relationship differed between years (Year by resistance level^2^ effect in ANOVA: F_1,47_ = 1.16, P = 0.28). However, when examining the relationship between resistance level^2^ and ASD separately between collection years, we found a significant negative quadratic relationship for the 2012 sample (solid line in Fig. 4; β = -0.48 ±0.23, F_1,22_ = 6.07, P = 0.02) but no evidence for a significant relationship among populations sampled in 2003 (β = -0.17 ±0.19, F_1,22_ = 1.08, P = 0.31). Despite the relative stability of ASD values among populations across this nine-year period (i.e., lack of a year effect), both floral traits comprising ASD – height of the tallest stamen and pistil height – showed a significant (or trend for significant) collection year by resistance level interaction (Pistil height: F_1, 47_ = 4.28, *P* = 0.04; Tallest stamen height: F_1, 47_ = 3.27, P = 0.08). In the 2012 sample, both the pistil and stamen height decreased as the level of resistance increased (2012: pistil β = -0.08 ±0.05, tallest stamen β = -0.08 ±0.05) whereas the pistil and stamen heights from the 2003 sample increased as resistance increased (2003: pistil β = 0.07 ±0.04, tallest stamen β = 0.06 ±0.05).

**Figure 4.**
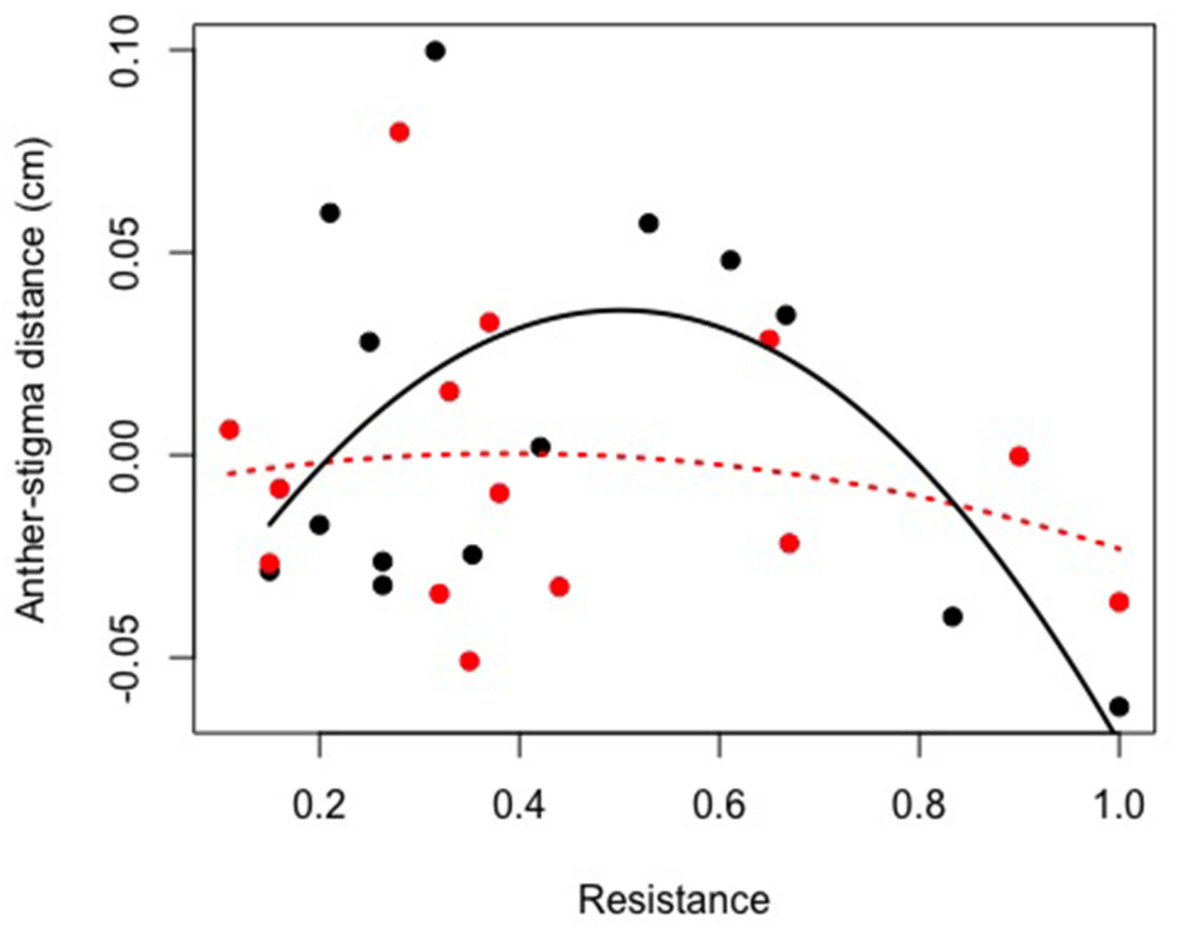
Anther-stigma distance (ASD) varies non-linearly with the level of resistance in 2012 (black dots; solid line) but not 2003 (red dots; dashed line). Fifteen populations sampled as seed in 2012 and 2003, respectively, were used for this analysis to determine if co-variation between ASD and glyphosate resistance varied between years. Shown are the residuals of anther-stigma distance (cm) according to resistance level at 1.7 kg ai/ha (averaged by population after removing variation due to sampling date). A significant quadratic relationship between ASD and resistance is present among populations sampled in 2012 (solid black line: β ± SE, 2012: -0.48±0.23; F_1,22_ = 6.07, P = 0.02) but not in 2003 (dashed red line: β± SE, 2003: -0.16±0.19; F_1,22_ = 1.08, P = 0.31).

## Discussion

In line with our predictions, we demonstrate that the mating system of the agricultural weed, *Ipomoea purpurea*, co-varies with the level of glyphosate resistance. Specifically, we find that outcrossing rates are lower and maternal inbreeding coefficients are higher in resistant compared to susceptible populations. We likewise find that anther-stigma distance, a floral trait associated with self-fertilization in this species, exhibits a nonlinear relationship with resistance such that the most and least resistant populations exhibit lower anther-stigma separation compared to populations with moderate levels of resistance. Further, this relationship was present among populations sampled in 2012 but not 2003, suggesting that reproductive traits may have rapidly evolved in these populations over the course of nine years. Below, we discuss each of our major findings and place them in the broad context of mating system changes associated with human-mediated selection.

## Plant mating changes associated with resistance

As anticipated by both the reproductive assurance (Baker 1955, 1974; Goodwillie *et al.* 2005; Pannell *et al.* 2015) and ‘prevention of gene flow’ (Antonovics 1968) hypotheses, our work finds a significant negative relationship between the outcrossing rate and level of herbicide resistance across natural populations of an agricultural weed indicating that individuals from highly resistant populations self more often than those from susceptible populations. We likewise uncovered a significant negative nonlinear relationship indicating that the outcrossing rate initially increased at low levels of resistance and then declined as resistance increased. When examining the nonlinear relationship using piecewise regression, however, we found that the initial positive slope was not significantly different than zero whereas the negative linear relationship following the estimated breakpoint was significant. Further, we found a positive, linear relationship between maternal inbreeding coefficients and resistance, indicating that maternal individuals from highly resistant populations were more likely to be the product of self-fertilization or mating between close relatives (*i.e*., biparental inbreeding) than maternal individuals from susceptible populations. These results overall show that the mating system is altered as populations increase in resistance level, indicating that the mating system may co-evolve with resistance.

There are currently few examinations of the mating system of xenobiotic tolerant or resistant species for comparison. In the most relevant example to date, metal-tolerant populations of the grass species *Anothoxanthum odoratum* and *Agrostis tenuis* exhibit higher self-fertility compared to nearby susceptible pasture populations (Antonovics 1968; Antonovics *et al.* 1971). Theoretical work by the same authors suggested that higher rates of selfing in the metal-tolerant populations should evolve to reduce the influx of non-adapted genotypes (*i.e*., the prevention of gene flow hypothesis) (Antonovics 1968); however, no potential mechanism was identified. Indeed, our findings of lower outcrossing in herbicide resistant populations along with a relationship between floral traits and resistance (*discussed below*) supports the prevention of gene flow hypothesis. Mechanisms that promote the self-pollination of adapted individuals would reduce the level of gene flow from non-adapted individuals (Levin 2010), thus ensuring that the offspring produced in novel or stressful/marginal environments are likewise stress-tolerant (Levin 2010). In this way, the increased self-fertilization of adapted types is hypothesized to lead to reproductive isolation between adapted and non-adapted individuals (Antonovics 1968; McNeilly & Antonovics 1968), and potentially promote niche differentiation (Levin 2010). Such shifts toward a higher propensity to self-fertilize in stressful habitats may not be unusual in nature, as a higher propensity to self has been identified in metal tolerant populations of *Armeria maritime* (Lefebvre 1970) and *Thlaspi caerulescens* (Dubois *et al.* 2003) as well as serpentine tolerant *Mimulus* (Macnair & Gardner 1998). We emphasize, however, that the above cases compare the ability to produce seed autonomously in greenhouse conditions between adapted and non-adapted individuals, which does not always correlate with selfing rates in nature, whereas our work presents estimates of the outcrossing rate of *I. purpurea* sampled from natural conditions.

Our data are likewise consistent with the reproductive assurance hypothesis. Originally proposed by Baker (1955), this hypothesis predicts greater selfing ability in colonizing species since they are likely mate-limited when arriving to new areas. Reproductive assurance through self-fertility is broadly, although not ubiquitously, supported by empirical research in other plant systems, such as small or pollinator-limited populations of *Capsella* (Foxe *et al.* 2009; Guo *et al.* 2009), *Leavanworthia* (Busch *et al.* 2011) and *Clarkia* (reviewed in Busch & Delph 2012). Agricultural weeds that experience selection from herbicide application or population reduction *via* other means, such as tilling, are analogous to species that colonize novel or new habitats. For example, in a scenario in which resistance alleles are at low frequency in the population and strong selection *via* herbicide application significantly reduces the population size, individuals that survive and re-colonize crop fields will then likely be mate limited. Resistant individuals with a higher propensity to self-pollinate would thus be at a relative advantage compared to those that cannot self-pollinate.

Another potential explanation for a relationship between higher selfing and resistance is the ‘segregation effect,’ wherein an allele that causes higher selfing enhances segregation and forms associations with homozygotes for beneficial (and other) alleles. As selfing modifiers become associated with the beneficial mutation, selfing individuals respond quickly to selection, which will then lead to even higher rates of selfing in the population (Uyenoyama & Waller 1991). In support of the segregation effect, recent multilocus simulations of the causes and consequences of selfing find a shift from outcrossing to high levels of selfing following the introduction of large-effect beneficial mutations, so long as the beneficial mutations have moderate to large fitness effects (Kamran-Disfani & Agrawal 2014). This dynamic is thus proposed for species that are establishing a new habitat, or following episodes of environmental change (Kamran-Disfani & Agrawal 2014) – both of which are experienced by plants exposed to herbicide application. Regardless of whether the data reported herein are best explained by selection for reproductive isolation (Antonovics 1968), reproductive assurance (Pannell *et al.* 2015), or the segregation effect (Kamran-Disfani & Agrawal 2014), the patterns we have uncovered among naturally-occurring populations of this common weed show that the plant mating system can be influenced by and evolve rapidly in response to selection following the application of herbicides.

## Anther-stigma distance co-varies with herbicide resistance

Initially, and in line with the ‘prevention of gene flow’ hypothesis, we predicted that anther-stigma distance should decrease as the level of herbicide resistance increases, since a lower anther-stigma distance leads to an increased rate of selfing in this species (Chang and Rausher 1997). Strikingly, we found a significant negative quadratic relationship between ASD and resistance but no evidence of a negative linear relationship across all populations as uncovered in the mating system data. Specifically, at very low levels of resistance (e.g. from 0 to 30% survival within the population), anther stigma distance increased with the level of resistance, but after 30% resistance, ASD decreased as resistance increased. One interpretation of this pattern is that at low levels of resistance, inbreeding depression in selfed progeny may lead to selection against low-ASD types whereas at higher levels of resistance, increased selfing due to low ASD confers a fitness advantage (i.e., reducing the influx of non-adapted alleles) that is greater than the cost associated with inbreeding.

An alternative explanation for the pattern between ASD and resistance is that low-ASD may be favored in both low and high resistant populations due to some other agent of selection that is potentially similar between the two types of populations. This explanation is less likely than the first, however, since low-ASD populations that are low- or high-resistance are both found in very different areas of the landscape – both R and S populations are from TN and NC/SC – where edaphic factors such as elevation and rainfall are very different (see Fig. 1). Further, although we found no evidence of a negative linear relationship between ASD and resistance across all sampled populations, the ASD of the least resistant populations was twice as large as the ASD from the most resistant populations (ASD, <20% resistant, N = 6: 1.00 ±0.10 (mm); >80% resistant, N = 4: 0.50 ±0.20 (mm)). Thus, despite the initial increase in ASD with resistance, the most resistant populations exhibit significantly lower ASD than the least resistant populations. As above, we note that for the mating system and associated floral traits to evolve in response to resistance evolution, the benefit of producing selfed progeny should outweigh any associated cost of inbreeding. While previous work has shown evidence of inbreeding depression in a single population of this species (Chang and Rausher 1999a), the level of inbreeding depression discovered was not strong enough to counteract the transmission advantage of selfing (i.e., delta < 0.50; Chang and Rausher 1999a). Further work will thus be required to determine if the costs of inbreeding relative to the benefits of selfing are responsible for the changes to the mating system and ASD that we describe herein.

Unlike reports from other weedy species (e.g. *Eichhornia paniculata,* Vallejo-Marín & Barrett 2009), we uncovered no evidence that decreases in ASD are the result of increased stamen length, and further, we found only marginal evidence that pistil height differences among populations may explain the pattern between ASD and resistance. Although we cannot identify which component of the composite trait ASD is responsible for the pattern uncovered with resistance, we do have evidence to suggest the relationship is more pronounced in the more recent population sampling (in 2012 versus 2003). While the majority of the populations sampled in 2003 had experienced glyphosate application (Kuester *et al.* 2015b), they further experienced consistent glyphosate application between sampling periods, and on average, populations in 2012 exhibit slightly higher levels of resistance compared to the same populations from 2003 (Kuester *et al.* 2015b). Interestingly, we uncovered a significant interaction between sampling year and resistance for pistil height: the relationship between pistil height and resistance was negative in 2012 but positive in 2003. This again suggests that perhaps the pistil length is co-evolving with resistance. However, further work targeting the nature of the correlation between these two traits will be required to allow us to draw this conclusion with certainty.

As hypothesized above, the responses to selection and subsequent changes to the mating system and associated floral traits that we have identified may be adaptive and due to a genetic basis, or could alternatively be due to plastic changes influenced by the environment (Rick *et al.* 1977; Brock & Weinig 2007; van Kleunen 2007; Vallejo-Marín & Barrett 2009). Several lines of evidence suggest that adaptation is the more likely cause for the patterns observed in this study. While we present field estimates of the mating system among populations, we measured floral morphology in a completely randomized greenhouse experiment using replicate maternal lines from many populations, with all individuals experiencing a common environment. Thus, the differences we report in ASD among populations very unlikely to be explained by different environmental factors from different populations. Further, both anther-stigma distance and glyphosate resistance have an additive genetic basis in this species (Chang & Rausher 1998; Baucom & Mauricio 2008; Debban *et al.* 2015), both traits respond rapidly to artificial selection (Chang & Rausher 1998; Debban *et al.* 2015), and populations sampled for the work reported herein were all from agricultural fields with a history of glyphosate application. Taken together, these data suggest that the co-variation we have uncovered between floral phenotypes and resistance across many natural populations of this weed is likely due to adaptive changes following selection rather than plasticity in either the mating system or the level of ASD.

## Future directions & conclusions

Although anecdotal reports suggest many herbicide resistant plants are predominantly selfing (Jasieniuk *et al.* 1996), ours is the first to identify co-variation between herbicide resistance and estimates of the outcrossing rate, thus providing empirical evidence that the mating or breeding system of a plant may co-evolve with resistance. We note, however, that the results we present cannot address the causal nature of the relationship between resistance and the mating system; although we discuss the dynamic as if the mating system co-evolves in response to the evolution of resistance, it is entirely possible that a highly selfing mating system is responsible for, or has maintained, the high levels of resistance within some populations. Further, like that described in the ‘segregation effect’ hypothesis, the evolution of the two traits may be intertwined such that resistance evolves first, following which selfing modifiers become linked to the beneficial resistance allele which then leads to higher rates of selfing evolving in the population. Future work will thus be required to disentangle the nature of the relationship between resistance and the mating system.

Overall, we have demonstrated that individuals from herbicide resistant populations self more than those from susceptible populations in natural settings and that low anther-stigma distance may be a potential mechanism underlying this increased rate of selfing. Our work identifies human impacts on plant mating patterns that go beyond the indirect consequences of environmental manipulations such as forest fragmentation and metal contamination. Changes in mating systems can have cascading effects on the effective population size (Nunney 1993), gene flow and the genetic diversity of natural populations (Hamrick & Godt 1996) and can determine the overall evolutionary propensity of these species. Our findings thus highlight the importance of considering the influence of human-mediated selection on correlated responses of natural populations that can lead to long-term evolutionary consequences. Likewise, these results show that associations between highly beneficial traits and plant reproduction can occur rapidly within ecological timescales. The results of our work are thus applicable to other scenarios of strong selection such as climate change or scenarios wherein individuals of colonized populations are mate-limited.

## Acknowledgements

We thank A Wilson for assistance and D. Alvando-Serrano, M. Van Etten, T-L Ashmann, V. Koelling, C. Dick, L. Moyle, J. Vandermeer, S.I. Wright, and S.C.H. Barrett for comments on earlier drafts of the manuscript. This work was funded by USDA NIFA grants 04180 and 07191. The authors declare no competing financial interests. Correspondence and requests for materials should be addressed to RSB. Primary data used in these analyses will be made available in the public github repository https://github.com/rsbaucom/MatingSystem2015, which can be anonymously accessed.

